# Model-based analysis of the stiffness of the wrist joint in active and passive conditions

**DOI:** 10.1101/281204

**Authors:** Andrea Zonnino, Fabrizio Sergi

**Author notes:** Address all correspondence related to ASME style format and figures to this author.

## Abstract

The control of joint stiffness is a fundamental mechanism used to control human movements. While many studies have observed how stiffness is controlled for tasks involving shoulder and elbow motion, a limited amount of knowledge is available for wrist movements, though the wrist plays a crucial role in fine manipulation.

We have developed a computational framework based on a realistic musculoskeletal model, which allows to calculate the passive and active components of the wrist joint stiffness. We first used the framework to validate the musculoskeletal model against experimental measurements of the passive wrist joint stiffness, and then to study the contribution of different muscle groups on the passive joint stiffness. We finally used the framework to study the effect of muscle co - contraction on the active joint stiffness.

The results show that thumb and finger muscles play a crucial role in determining the passive wrist joint stiff - ness: in the neutral posture, the direction of maximum stiffness aligns with the experimental measurements, and the magnitude increases by 113% when they are included. Moreover, the analysis of the controllability of joint stiffness showed that muscle co - contraction positively correlates with the stiffness magnitude and negatively correlates with the variability of the stiffness orientation (p < 0.01 in both cases). Finally, an exhaustive search showed that with appropriate selection of a muscle activation strategy, the joint stiffness orientation can be arbitrarily modulated. This observation suggests the absence of biomechanical constraints on the controllability of the orientation of the wrist joint stiffness.

## 1 Introduction

Object manipulation is a crucial human activity that involves physical interaction between the human body and an external environment. It is one of the most complex task for the neuromuscular system since, to be successfully completed, it requires simultaneous control of the movement accuracy and of the stability margins. Some tasks may, in fact, be intrinsically unstable and tend to deviate the limb from the desired movement [1].

The Central Nervous System (CNS) of healthy humans is capable of tuning the dynamic response of the musculoskeletal system during interactions with the environment, thereby compensating for possible instabilities, and producing smooth and graceful movements [2–6]. A paramount mechanism adopted by the CNS to control for the stability margins during physical interaction with the environment is modulation of joint stiffness using muscle co - contraction [7, 8]. Since for a muscle undergoing isometric contractions the number of actin - myosin cross - bridges increases with muscle force, and since a greater number of cross - bridges results in greater muscle stiffness, skeletal muscles have force - dependent stiffness properties. [9,10]. As such, by appropriate modulation of the forces applied by an agonist - antagonist set of muscles, it is possible to modulate the joint stiffness independently from joint torque. Greater values of stiffness afford greater stability margins when the musculoskeletal system is perturbed.

Because of the nature of the muscle tissues, the joint stiffness depends on the intrinsic passive - elastic properties of the muscles within the arm, i.e. passive component, in addition to the their activation level, i.e. active component. Experimental studies attempted to characterize both components for a single - joint and multi - joint system by asking individuals to relax their muscles [11–13] — to measure the passive component — or to apply isometric forces at different levels of co - contraction [5, 14–16] — to measure the active one. In these studies, a robotic manipulator was used to apply displacements to the human joint and measured the resistive torque. Although these studies were able to investigate stiffness at the joint, or end - effector, level, there are no studies that investigated how different muscles groups affect the compound value of joint stiffness in the active and passive case.

Another crucial and poorly understood point refers to the constraints on the achievable joint stiffness values. While for a simple 1 degree of freedom (DOF) joint, co - contraction of agonist - antagonist muscle groups is undoubtedly recognized as the main mechanism used to modulate the joint stiffness magnitude [17–21], much less is known for more complex multi - DOF systems. In these systems, the multi - dimensional nature of the task space opens the possibility of independent modulation of stiffness magnitude and orientation. So far, controversial results have been obtained for movements involving combined motion of the shoulder and of the elbow. One set of experiments studying planar reaching movements found that, after some training, the CNS can almost arbitrarily regulate both the magnitude and orientation of the end - effector stiffness without constraints [22–25]. On the contrary, another set of experiments based on isometric forces concluded that the CNS cannot arbitrary regulate such properties, and that it is only able to modify the orientation of the end - effector stiffness by roughly 30 deg [26–28]. The latter results suggest that the neuromuscular system might have neural and / or biomechanical constraints that limit the range of achievable joint stiffness values; however this is only a speculation based on evidence presented so far. To properly rule out whether such constraints arise from a neural or biomechanical origin, it would be important to search exhaustively for the set of muscle co - contraction patterns admissible for a given task, and derive the entire set of achievable stiffness values: if the variability of the achievable behavior (stiffness magnitude and orientation) is large, then the constraint ought to be neural; if the behavior is of limited variability, then the constraint ought to be primarily biomechanical.

Most of the analyses on the value of joint stiffness and its controllability have focused on the shoulder - elbow group. However, being the wrist the most distal joint of the upper arm, it has a key role in controlling the dynamics of fine manipulation tasks. Recently, some studies have been done to investigate the dynamics of the wrist rotations [29] and to estimate the 2D passive component of the joint stiffness in its neutral posture [11, 12]. Not much, however, is known about the controllability of the wrist stiffness with muscle co - contraction. Furthermore, even though the different muscles that actuate the fingers are highly coupled with the wrist joint [30], it is currently unknown how much different muscle groups affect the overall stiffness of the wrist joint.

In summary, despite the importance of studying of the wrist joint stiffness to understand how the CNS controls fine manipulation tasks, investigation of the joint stiffness has been done focusing mostly on the shoulder - elbow group. Thus, the capability of the CNS to control wrist stiffness and the effects of different muscle groups on the passive and active wrist stiffness have never been characterized. Performing these characterizations with *in - vivo* experimental procedures is impractical because of the impossibility to reliably separate the effect of many muscle groups on the compound joint stiffness. The use of musculoskeletal models (MSMs) is a promising approach to study these properties of wrist stiffness. MSMs have already been used to investigate the joint stiffness achievable during isometric force generation [31, 32]. Moreover, a few computational studies expanded those analyses to identify the contribution of different muscles in modulating joint stiffness for the shoulder and elbow group [33–35]; however, in these cases, simplified geometric and muscle models were used.

In this study, we present a novel computational framework based on a state - of - the - art musculoskeletal model imple - mented in OpenSim [36], which allows to analyze the wrist joint stiffness in terms of both passive and active components. Because our approach is based on a realistic musculoskeletal model, our analysis validated for the first time previous experimental measurements of wrist joint stiffness, and extended these measurements to multiple wrist postures. Moreover, via the use of virtual models that include only select muscle groups, we established the role of different muscle groups in determining the passive and active joint stiffness. Finally, our analysis based on the exhaustive search of muscle co - contraction patterns admissible for a given joint position and torque indicated that wrist joint stiffness can be controlled in a wide range of magnitudes and orientations.

## 2 Materials and Methods

### 2.1 Model

Our work is based on the MSM presented in [36] (Fig. 1, left). The model describes the geometry and mechanical properties of upper extremity muscles, tendons, and bones. For this work, we focused on four DOFs and 24 muscle segments (Tab. 1), those directly affecting the wrist joint. All properties of the model were evaluated for different values of the two DOFs of the wrist joint, i.e. Flexion/Extension (FE) and Radial / Ulnar Deviation (RUD), within a rectangular domain. The two remaining DOFs — forearm Prono / Supination (PS) and elbow Flexion / Extension (eFE) — were set to θ_*PS*_ = 0 deg, θ_*eFE*_ = 90 deg and kept constant, defining the limb’s posture. The hand posture was unchanged from the original model, and kept in a cylindrical grasp configuration, a posture commonly used to determine the passive stiffness of the wrist joint [11,12].

**Fig. 1.**
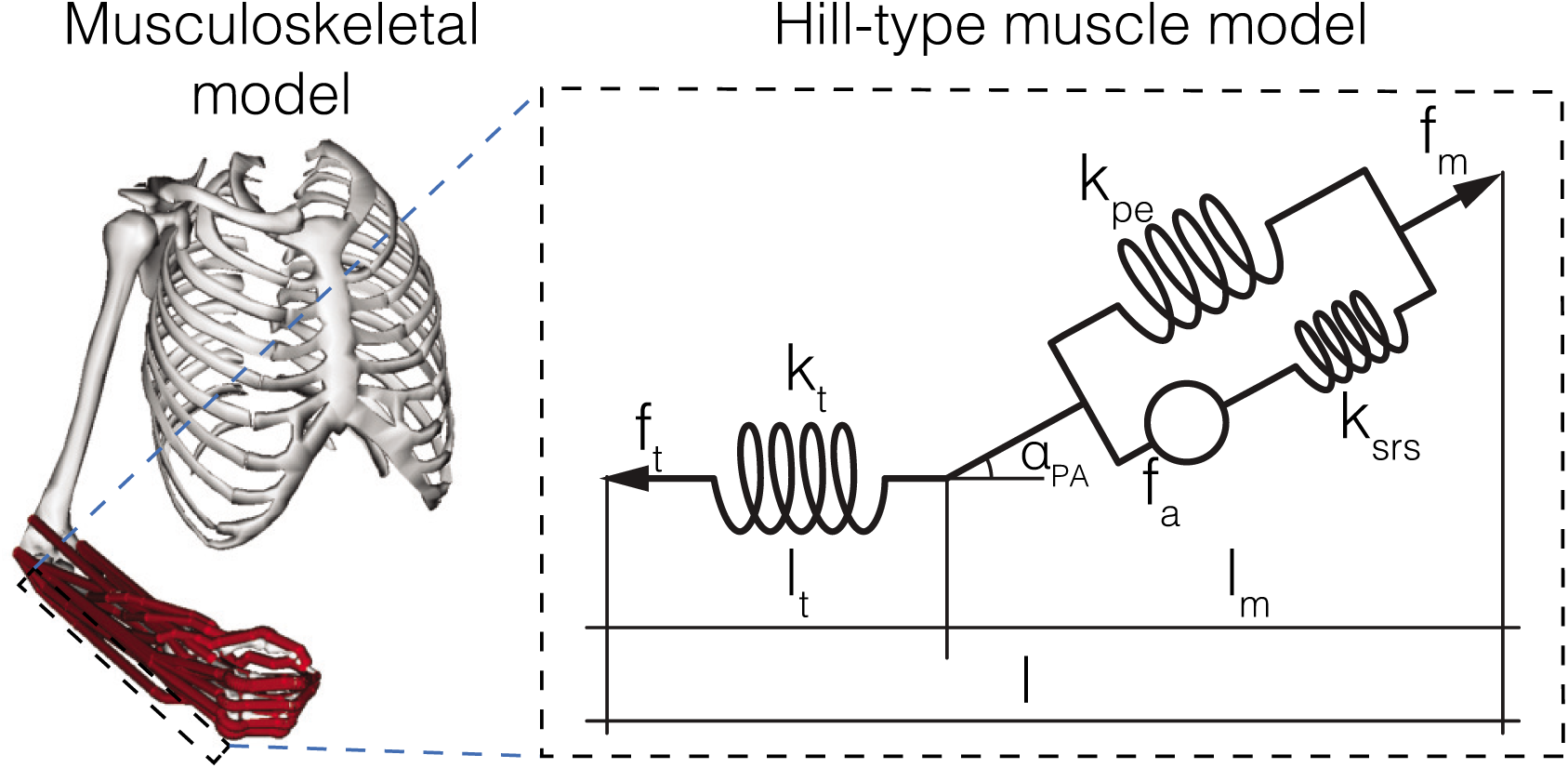
Representation of the musculoskeletal system (left) and the musculotendon unit (right) used for the analysis.

**Table 1.** Muscle groups in the model

| Group | Muscle Name | Abbreviation |
| --- | --- | --- |
| **Wrist** | Extensor Carpi Radialis Brevis | ECRB |
|  | Extensor Carpi Radialis Longus | ECRL |
|  | Extensor Carpi Ulnaris | ECU |
|  | Flexor Carpi Radialis | FCR |
|  | Flexor Carpi Ulnaris | FCU |
| **Palm** | Palmaris Longus | PL |
| **Thumb** | Extensor Pollicis Brevis | EPB |
|  | Extensor Pollicis Longus | EPL |
|  | Abductor Pollicis Longus | APL |
|  | Flexor Pollicis Longus | FPL |
| **Finger** | Extensor Digitorum Communis* | EDC |
|  | Extensor Indicis Proprius | EIP |
|  | Extensor Digiti Minimi | EDM |
|  | Flexor Digitorum Superficialis* | FDS |
|  | Flexor Digitorum Profundus* | FDP |

\* muscle composed of different segments, one per each finger: Index (I), Middle (M), Ring (R), Little (L). Since the model does not describe individual finger motion, this muscle is considered effectively as one muscle in this work.

We developed a framework to estimate the stiffness of the wrist joint in different postures defined by the wrist FE (θ_*FE*_) and RUD (θ_*RUD*_) coordinates. To estimate joint stiffness using the MSM, we used OpenSim to solve for the kinematics of the musculotendon (MT) units, thus obtaining muscle moment arm **r** (matrix that contains the moment arm of all muscles about all joint axes) and the lengths of the MT units **l**_*mt*_ [37, 38]. The bold notation is used for vectors and matrices.

The MSM includes a Hill - type model for each muscle [39, 40] (Fig. 1, right), with an active force generator *f*_*a*_ and two compliant elements, one in series to the force generator, *k*_*srs*_, and one in parallel to the series of *f*_*a*_ and *k*_*srs*_, *k*_*pe*_. This three - element muscle unit is then in series with a tendon, modeled with a passive elastic element, *k*_*t*_.

Given a value of activation coefficient a, muscle mechanics is completely determined by the set of MT coefficients: maximum isometric muscle force *F*_*MAX*_, active and passive force multipliers 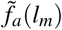 and 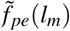, optimal muscle length *l*_0_, muscle pennation angle α_*PA*_(*l*_*m*_), muscle passive stiffness *k*_*pe*_, and tendon parameters such as tendon slack length *l*_*s*_, and tendon stiffness *k*_*t*_ (*l*_*t*_). As such, given *l*_*mt*_ we estimated the length of the muscle *l*_*m*_ and tendon *l*_*t*_ unit by imposing static equilibrium of the system, and then calculated the MT force as:

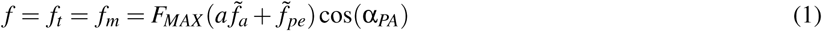

To enable the study of both active and passive muscle stiffness, we used the assigned passive force vs. length function 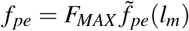 provided by OpenSim to estimate values of *k*_*pe*_ for different muscle length *l*_*m*_. This function describes the behavior of the muscle unit in absence of active force generation (*a* = 0), i.e. the passive component of the stiffness. The series compliant element *k*_*srs*_ describes instead the muscle’s capability of modulating stiffness with neural activation, i.e. the active component of the stiffness. The active component is commonly modeled with the Short - Range Stiffness (SRS), a model used to describe the initial response of a muscle when perturbations of small amplitude are applied [9, 10, 41, 42]. We based our SRS model on the one presented by [41], but modified it to highlight the fact that active stiffness increases only with the active force 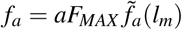, and not with total force that could result also from passive muscle stretching. As such, the SRS of each muscle was then calculated as:

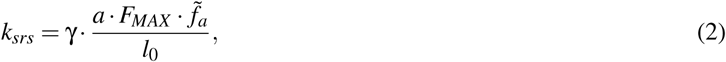

where γ is a dimensionless scaling factor previously measured in feline hind - limb muscles (γ = 23.4) [41], and indirectly validated in human subject [32]. In this work, we will use this value to determine the effect of muscle co - contraction on the modulation of muscle stiffness.

### 2.2 Mathematical derivation of the joint stiffness

In a system composed by *N* joint angles and *M* muscles, the joint torques are given by the following equation,

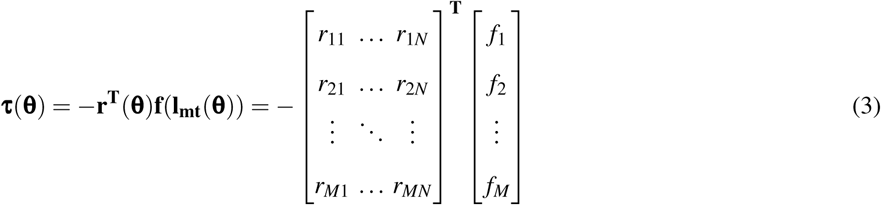

where **τ** is a *N* × 1 vector that contains values of torque for the *N* joint angles in the system, the component *r*_*ij*_ is the moment arm of the muscle *i* with respect of the joint angle *j*, and *f*_*i*_ is the MT force for the muscle *i* computed using Eqn. (1).

Joint stiffness is defined as:

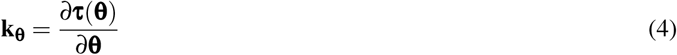

and can be calculated by substitution of Eqn. (1) and Eqn. (3) in Eqn. (4), and computation of the derivatives as:

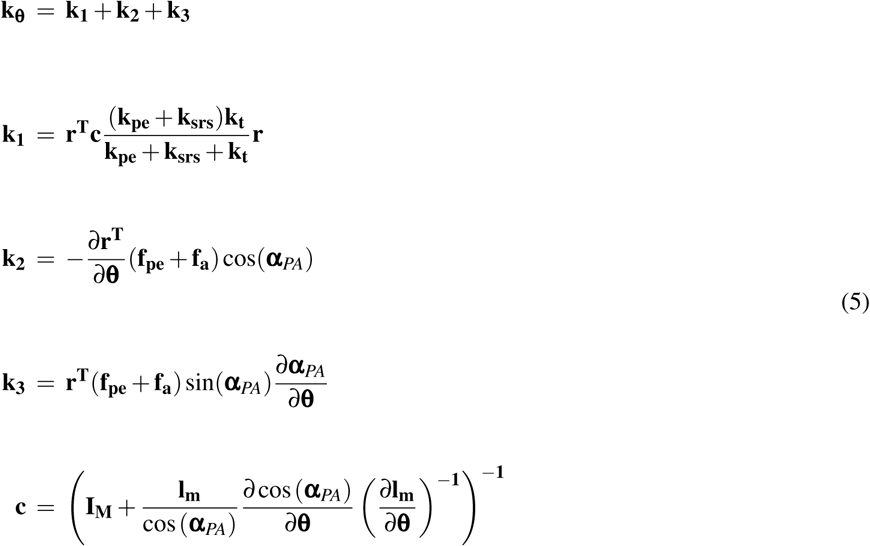

where muscle stiffness parameters have been converted into *M × M* diagonal matrices, and the remaining muscle parameters have been converted into *M* × 1 vectors containing coefficients for each muscle. Matrix **I_M_** is the *M × M* identity matrix. The derivative of the moment arm matrix with respect to the joint angle vector can be computed numerically as presented in [15].

Term **k_1_** describes how much a joint displacement changes the MT length thus stretching the spring - like elements in muscles and tendons. The second (**k_2_**) and third (**k_3_**) terms account for the joint - dependent change in muscle moment arms and pennation angles and how they affect joint stiffness. Even though **k_1_** is a quadratic form and always results in a positive definite contribution to joint stiffness **k_θ_**, **k_2_** and **k_3_** can have either positive or negative eigenvalues. Given our formulation, a negative eigenvalue of the joint stiffness matrix **k_θ_** corresponds to an unstable configuration, whereby for an applied displacement along a direction corresponding to the negative eigenvalue, the resulting joint torque is in the same direction of the displacement, causing the system to amplify the disturbance and quickly diverge from the initial posture. This is in contrast to what is expected from a stable spring - like behavior, where the force resulting from a perturbation opposes the applied perturbation [43]. **k_2_** in particular, known as the geometric stiffness because it includes the derivatives of the moment arms with respect of the joint angles, has been observed to cause possible instability also in a simple 1 - DOF model [44].

Equation (5) extends the formulation previously derived in [15] and accounts for the effect of the pennation angle and its variability across the workspace. Moreover, since our framework distinguishes the effect of passive and active muscle stiffness, it allows to separate the effect of such components also on the resulting joint stiffness.

### 2.3 Framework for joint stiffness calculation

The framework for the calculation of joint stiffness is graphically represented in the flowcharts in Fig. 2, which highlight the difference between passive and active muscle conditions.

**Fig. 2.**
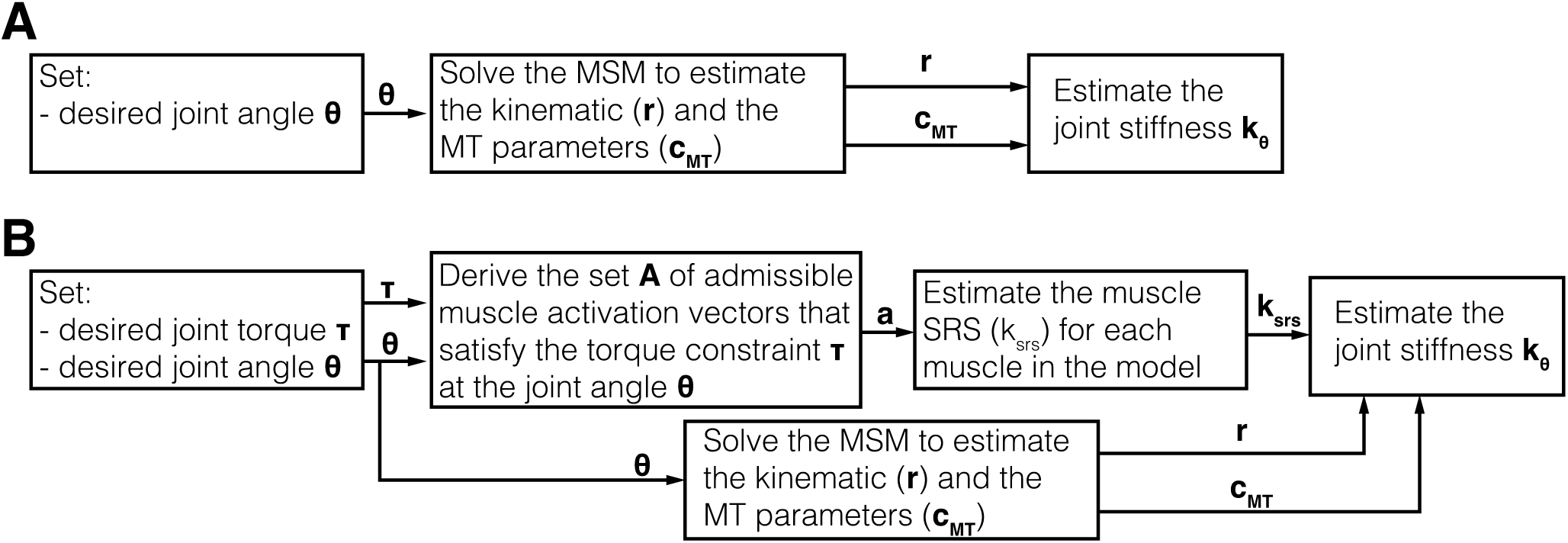
Block diagram of the framework used to calculate the passive (A) and active (B) joint stiffness.

In the passive case, given the joint angle vector **θ**, MSM kinematics is solved to derive the set of kinematic and MT parameters. These parameters are used in Eqn. (1) to calculate the passive muscle force; Eqn. (5) is then used to estimate joint stiffness. In these conditions, muscle force results uniquely from the stretching of passive fibers, so the net joint torque cannot be controlled and a single value of joint stiffness is calculated for any given posture.

In the active case, it is possible to modulate the net torque at the joint level by controlling the activation of the different muscles. Because of the redundancy of the musculoskeletal system, where the number of muscles is bigger than the number of DOFs, infinite activation vectors yield the desired joint torque for a given joint angle. To estimate the set of admissible joint stiffness values achievable for given joint position and torque constraint, the computational framework derives the set **A** ∈ ℝ^*M*x*W*^ of *W* admissible muscle activation vectors **a** ∈ ℝ^*M*^ that satisfy the joint torque constraints within a specified tolerance (10^*-3*^ tolerance). For every activation vector **a** within the admissible set, we compute muscle force using Eqn. (1), muscle SRS using Eqn. (2), and finally we calculate joint stiffness using Eqn. (5). This process leads to a set of joint stiffness values **K_θ_**∈ ℝ^*NxN*x*W*^ for every value of desired joint torque and joint angle.

The stiffness of a 2 - DOF system is commonly represented as an ellipse [5, 45]. The length of the two semi - axes quantifies the force opposed by the system when a unit displacement is applied along the semi - axes directions. In general, the distance between the center of the ellipse O and a point on the ellipse P quantifies the magnitude of the force in response to a perturbation along the direction of vector OP. Using the stiffness - ellipse analogy, we characterize each stiffness matrix via the following metrics:

- Magnitude *H*, calculated as the geometric mean of the eigenvalues *r*_*MAX*_ and *r*_*MIN*_ of matrix **k**_θ_, *H* ∈ [0, ∞) N·m/deg;

- Orientation Φ, defined as the relative angle of the principal axis of the ellipse with the horizontal axis, Φ ∈ (- 90, 90] deg;

- Eccentricity *e*, defined as 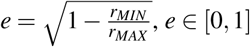

In the following sections, we report on the use of the developed framework first to extend experimental studies aimed at characterizing passive joint stiffness, and then to study the effect of different muscle groups on the stiffness of the wrist joint.

### 2.4 Passive stiffness analysis

#### 2.4.1 Validation

We first validated the framework against previous experiments that measured the passive wrist joint stiffness [11, 12]. Given limitations in the experimental measurement of joint stiffness, with available techniques requiring perturbations about multiple directions around a single posture, there is availability of experimental data only in the neutral posture, which corresponds to coordinates **θ** = [0, 0] deg of our model.

We quantified the mismatch between the estimate of passive wrist joint stiffness **k_θ_** obtained with our framework and the two values reported by [11] and [12] by calculating the difference in the three selected stiffness metrics (*H*, Φ, and *e*).

#### 2.4.2 Effect of different muscle groups

We then sought to isolate the contribution of different muscle groups to the passive wrist joint stiffness. To this aim, we divided the muscles shown in Tab. 1 in the following groups:

1. Wrist muscles: ECRL, ECRB, ECU, FCR, FCU;
2. Palm muscle: PL
3. Thumb muscles: EPL, EPB, APL, FPL
4. Finger muscles (I): EDC, FDS, FDP
5. Finger muscles (II): EIP, EDM

and created five virtual models that include different combinations of these groups: (PM1) only wrist muscles; (PM2) wrist and palm muscles; (PM3) wrist, palm and thumb muscles; (PM4) wrist, palm, thumb and finger (I) muscles; (PM5) wrist, hand, thumb and finger (I and II) muscles. For each virtual model, we computed the passive wrist joint stiffness in the neutral posture, along with the three metrics *H*, Φ, and *e*.

To quantify the effect of different muscle groups on the passive wrist joint stiffness, we calculated the difference in the stiffness metrics between each virtual model and the values obtained via human subject experiments [11] and [12].

#### 2.4.3 Passive stiffness map

We finally used the framework to extend previous experimental measurements and estimate the passive stiffness of the wrist joint in a range of FE and RUD angles defined by θ_FE_ *×* θ_RUD_ = [-40, 40] *×* [-5, 15] deg. Values were selected by discretizing the rectangular grid with spacing Δ_*FE*_ = 5 deg and Δ_*RUD*_ = 2.5 deg. For each posture, the joint stiffness matrix **k_θ_** along with the related metrics *H*, Φ, and *e*, were calculated.

To establish the variability introduced by different postures on the passive joint stiffness, we calculated the coefficient of variation *c*_*v*_ as the ratio between the standard deviation and the mean of the extracted distribution for each of the three stiffness metrics. Since the orientation is an axial variable, mean and standard deviation of this metric have been computed according to [46].

### 2.5 Active Stiffness

#### 2.5.1 Effect of co - contraction

We quantified how muscle co - contraction modulates joint stiffness in the wrist’s neutral posture by determining the set of admissible joint stiffness values for a zero net joint torque applied. Given the high degree of redundancy of this system (*M* = 15, *n* = 2, giving a 13 - dimensional manifold of activation vectors that satisfy the joint torque constraint for a given posture) we discretized the activation of *M - n* muscles in increments of 25%, yielding 5 possible activation values per muscle, and calculated all combinations of activation values for the *M -n* muscles, yielding 5^13^ possible activation vectors. Activation values were predefined for all muscles with the exclusion of ECRL and FCU; their activation, *a*_*i*_ and *a* _*j*_, were determined by solving the following equation:

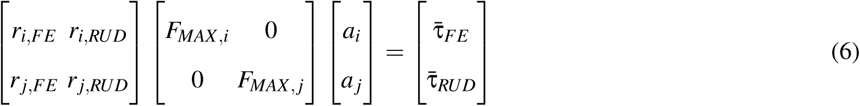

where 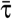 includes the torque resulting from all passive muscles, and from the active force of the muscles with predefined activation.

If any of the activation coefficients obtained via Eqn. (6) was outside the [0,1] range, then the calculated activation vector was considered unfeasible and discarded. The set **A**^(0)^ was then populated by all feasible activation vectors **a** calculated via this process. For every feasible activation vector **a**, the level of muscle co - contraction was quantified by the Global Activation Level (GAL), defined as:

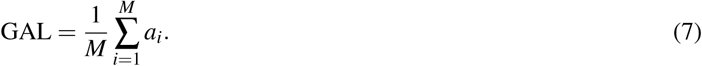

Given the feasible set of activations **A**^(0)^, the framework was applied to calculate the set of stiffness values 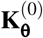 admissible for the given posture and joint torque, quantified in terms of metrics *H*, Φ, and *e*.

We then quantified the variability of the resulting stiffness metrics by measuring the coefficient of variation of the three stiffness metrics. We assessed the relationship between the magnitude of the achievable set of joint stiffness values and the level of muscle co - contraction using a linear regression analysis between the metrics *H* and GAL.

We finally assessed the relationship between GAL and the capability of controlling the orientation of the stiffness ellipse by grouping the measured ellipses in bins of 10% GAL increments, and calculating the mean and standard deviation of the stiffness orientation Φ in the different bins. The presence of a significant association between level of muscle co - contraction and the stiffness orientation and variability in each bin was established via two separate linear regressions between the mean and standard deviation of Φ within each bin and an ordinal variable (group I, group II, etc.) representing GAL. In all statistical testing, significance was set at the 0.05 type I error rate.

#### 2.5.2 Effect of different muscle groups

We finally investigated the effect of co - contraction of different muscle groups on the achievable set of joint stiffness values. We divided the muscles of the MSM in two groups; then, by using the same method described above, we estimated the achievable wrist joint stiffness sets when each muscle group was allowed to co - contract, separately. When only one muscle group was allowed to co - contract, the remaining muscles were kept in a passive condition. The two groups were defined as:

(AG1) Wrist and palm muscles, composed of ECRL, ECRB, ECU, FCR, FCU, PL;

(AG2) Thumb and finger muscles, composed of EPL, EPB, APL, FPL, EDC, FDS, FDP, EIP, EDM

Given the smaller dimensionality of these two problems (i.e. *M -n* = 4 for AG1, *M -n* = 7 for AG2) compared to the one of the complete model, we could derive the sets of admissible activation vectors **A** in a finer search space using exhaustive search. For AG1, we discretized the activation of *M -n* muscles in increments of 5%, yielding 21 possible activations values per muscle, and calculated all combinations of activation values for the *M - n* muscles, yielding 21^4^ activation vectors. We ran the systematic search for ECRB, ECU, FCR, and PL, and for each tested activation vector, we calculated the activation of ECRL and FCU compatible with the joint constraint using Eqn. (6). Similarly, we repeated the procedure for AG2, using this time a 10% increment in muscle activation values to obtain 11^7^ possible set of activation vectors. We ran the systematic search for all muscles in the group with the exception of FDS and EDC, and then calculated the activation of FDS and EDC compatible with the joint torque constraint. With these procedures, we derived the sets of muscle activations **A**^(AG1)^ and **A**^(AG2)^, corresponding to the search based on AG1 and AG2 muscles respectively, which led to the calculation of the set of achievable joint stiffness values for each muscle group, **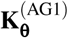** and **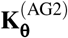** respectively.

Once obtained the sets of achievable joint stiffness, we quantified their variability by measuring mean, standard deviation, and coefficient of variation, separately for the two muscle groups, using the metrics *H*, Φ, and *e* defined above. To compare the ability of different muscle groups to modulate joint stiffness, we conducted pairwise comparisons of the estimated metrics between the two calculated sets using *t* tests.

## 3 Results

### 3.1 Passive stiffness

#### 3.1.1 Validation

Comparison between the physiological values of passive wrist joint stiffness and the ones obtained with model estimate shows errors in orientation (3 deg difference with [11] and 12 deg difference with [12]) and eccentricity (1% difference with [11] and 23% difference with [12]) (Fig. 3, Tab. 2, model PM5) that can be considered small. On the contrary, the magnitude of the MSM estimate is 58% lower than the value measured by [11], and of 47% lower than the value measured by [12].

**Fig. 3.**
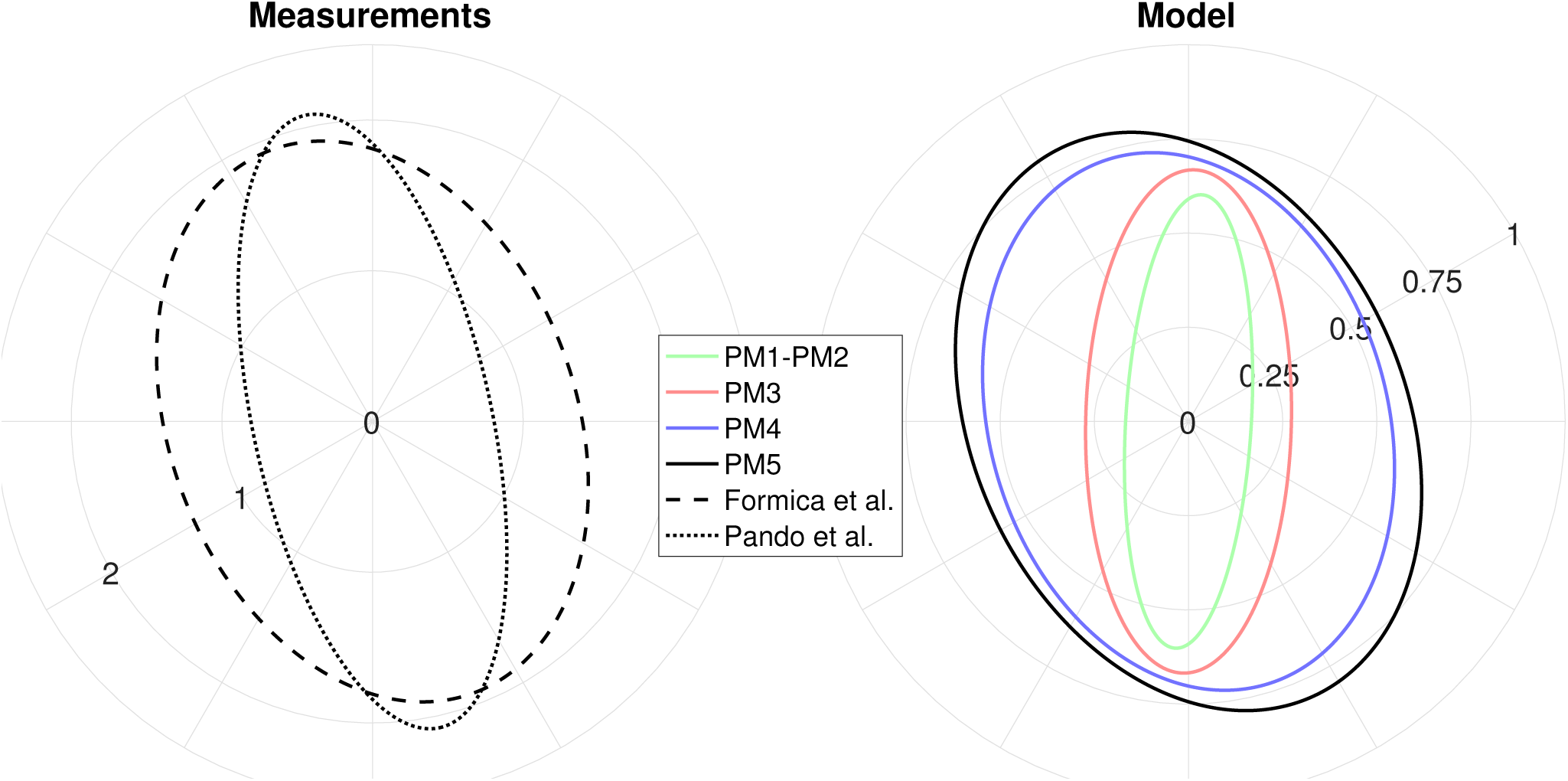
Measurements of the passive stiffness of the wrist (left). Model - based estimates of the passive stiffness when different groups of muscles are included (right). All values are obtained with the wrist in the neutral posture. Units of stiffness in the polar plot are N m deg^-1^.

**Table 2.** Contribution of the different muscle groups to the metrics that quantify joint stiffness

| Groups | Magnitude [N·m/deg] | Orientation [deg] | Eccentricity [-] |
| --- | --- | --- | --- |
| **Measured\*** | 1.61 | -69.27 | 0.71 |
| **Measured\*\*** | 1.28 | -77.61 | 0.92 |
| PM1 | 0.32 | 86.64 | 0.96 |
| PM2 | 0.32 | 86.64 | 0.96 |
| PM3 | 0.43 | 88.74 | 0.91 |
| PM4 | 0.62 | -73.38 | 0.69 |
| PM5 | 0.68 | -65.71 | 0.70 |

#### 3.1.2 Effect of different muscle groups

The progressive inclusion of the palm, thumb and finger muscles has effects on all considered stiffness metrics (Tab. 2, Fig. 3). Specifically, the direction of maximum stiffness is rotated toward the experimental measurements reported by [11, 12], the magnitude is increased — the complete model PM5 has a stiffness magnitude 113% greater than model PM1 — and the ellipse’s eccentricity is decreased toward the values reported in [11].

#### 3.1.3 Passive stiffness map

The passive wrist joint stiffness map is shown in Fig. 4. Analysis of the three stiffness metrics showed a mean *±* standard deviation of *H* = 0.69 *±* 0.28 N·m / deg for the magnitude, of Φ = - 58.96 *±* 24.38 deg for the orientation, and of *e* = 0.81 *±* 0.38 for the eccentricity. As a consequence, the coefficients of variation for the three metrics are *c*_*v,H*_ = 0.39, *c*_*v*,Φ_ = 0.20, *c*_*v,e*_ = 0.47. The distribution of the stiffness orientation is shown in Fig. 5 in a polar histogram plot, which shows the bi - modal nature of this distribution, with peaks around - 60 deg and 90 deg.

**Fig. 4.**
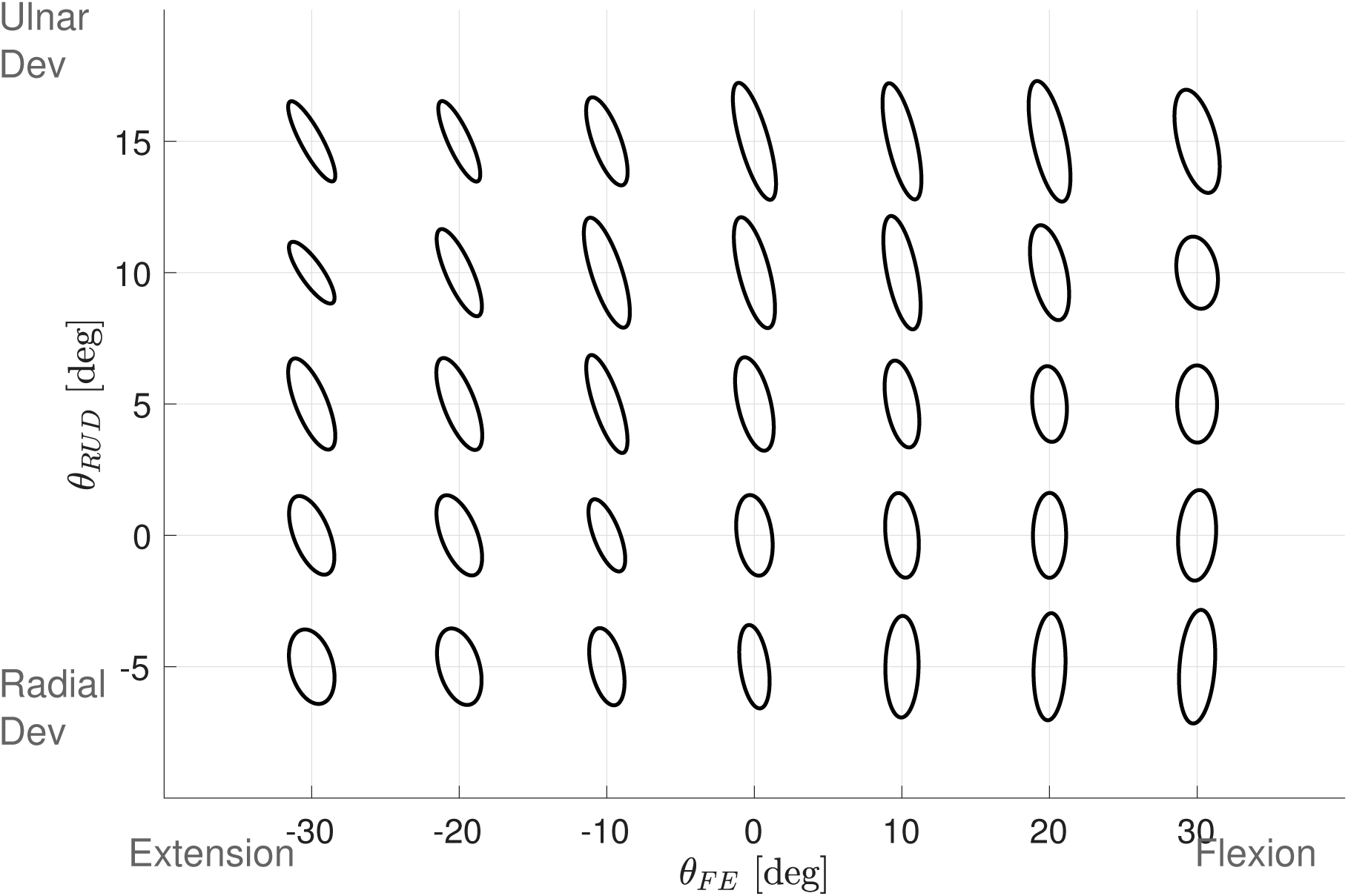
Map of the passive stiffness of the wrist joint in multiple postures.

**Fig. 5.**
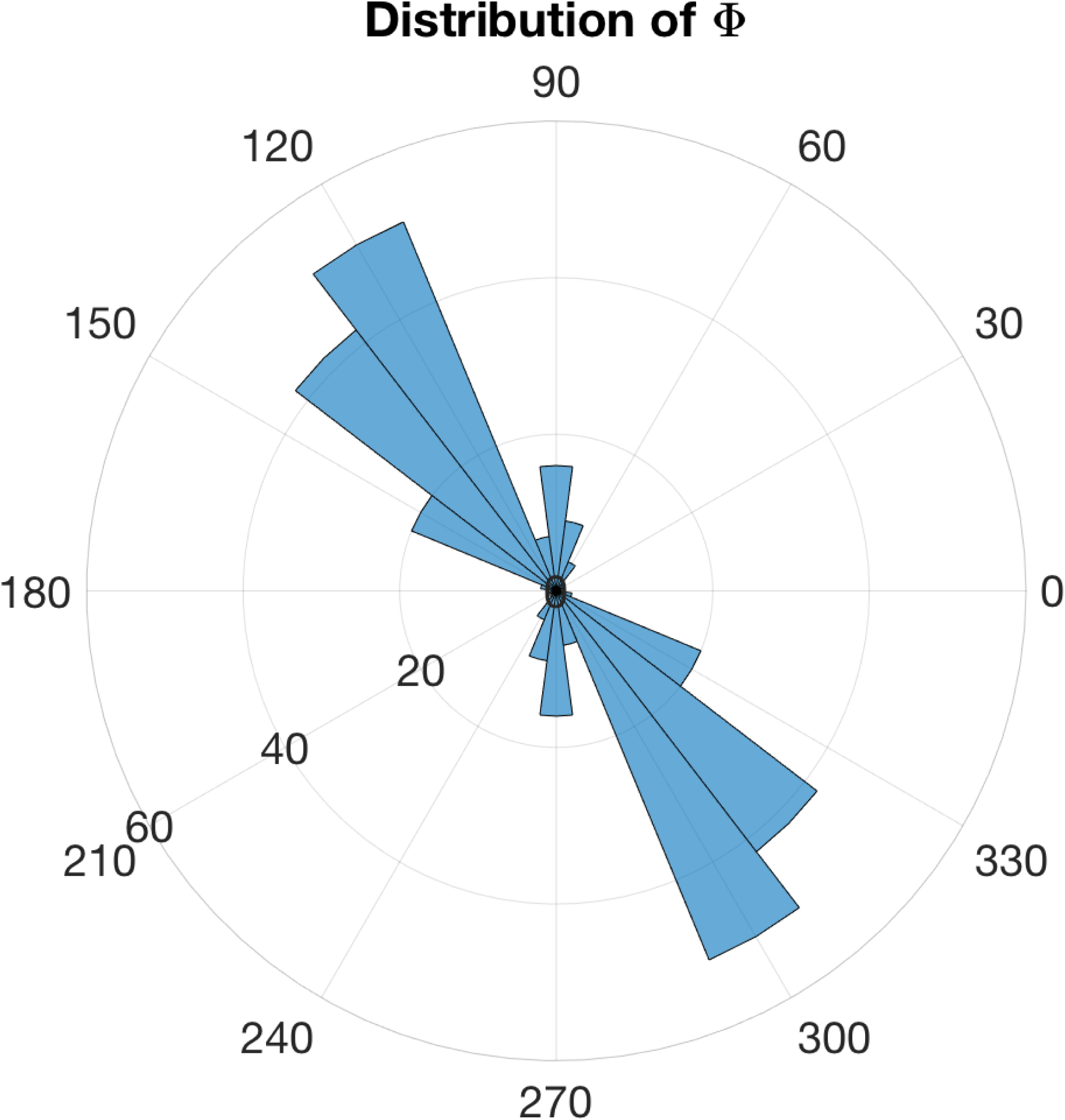
Polar histogram representing the distribution of the orientation of the passive stiffness of the wrist joint in the joint angle domain considered.

### 3.2 Active stiffness

#### 3.2.1 Effect of co - contraction

The systematic search resulted in 12,703,892 compatible activation vectors. That corresponded to 1% of the tested activation vectors. The achievable set of joint stiffness **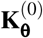** is represented in Fig. 6 (left), with the different ellipses color - coded by GAL. The analysis of the three stiffness metrics showed a mean *±* standard deviation of *H* = 28.68 *±* 4.74 N·m / deg for the magnitude, of Φ = 59.98 *±* 17.44 deg for the orientation, and of *e* = 0.83 *±* 0.10 for the eccentricity, with corresponding coefficients of variation *c*_*v,H*_ = 0.16, *c*_*v*,Φ_ = 0.29, *c*_*v,e*_ = 0.08.

**Fig. 6.**
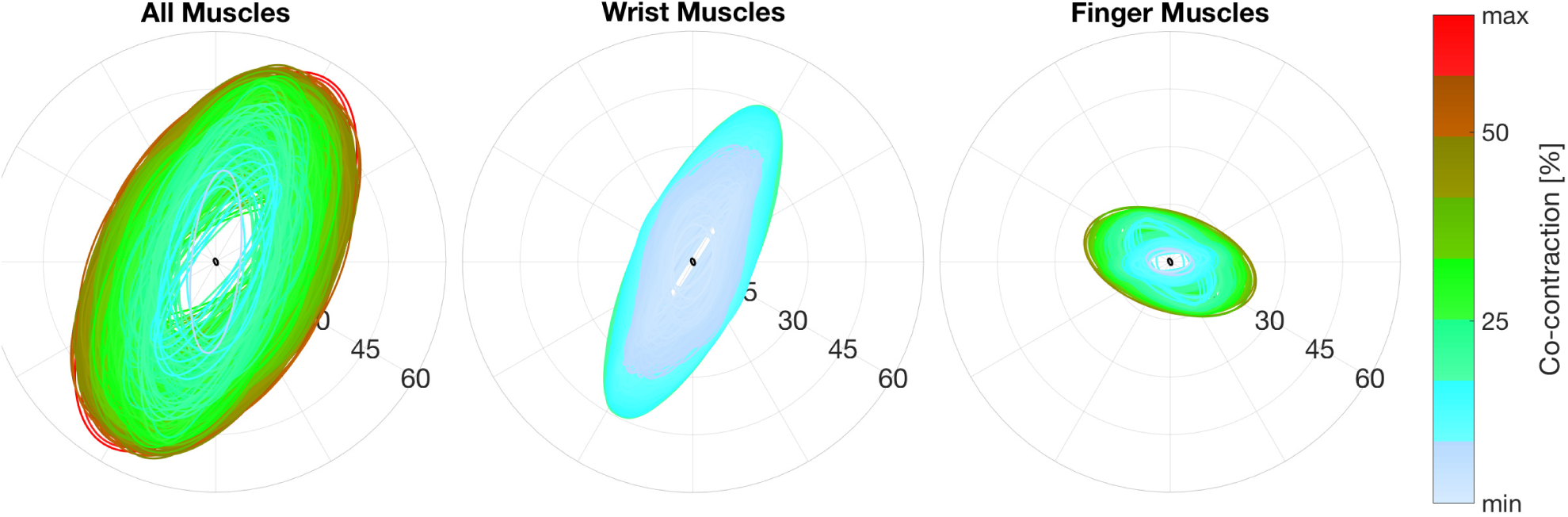
Set of admissible stiffness values established via systematic search when all muscles (left) can co - contract, or when only the wrist muscles (center), or only the finger and thumb muscles (right) can co - contract. The values refer to to the neutral posture and to a null desired net torque. The black ellipse shows the value of passive stiffness. Units of stiffness in the polar plot are N m deg^-1^.

**Table 3.**
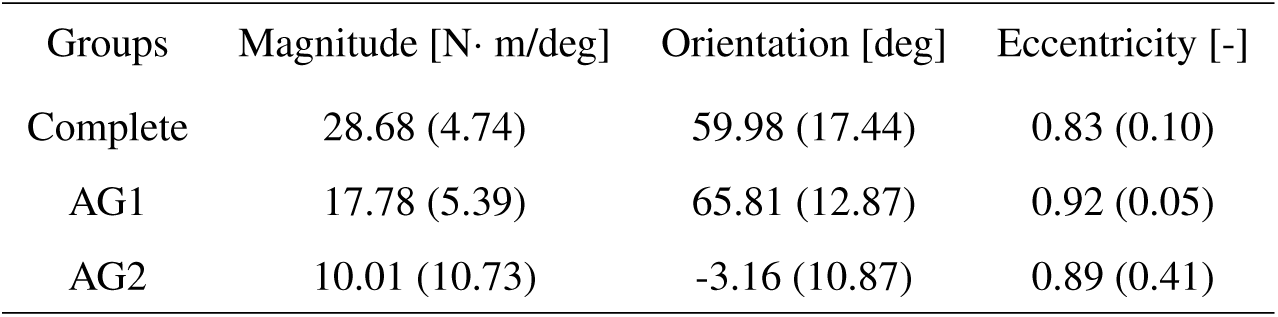
Contribution of the different muscle group to the metrics that describe the joint stiffness

| Groups | Magnitude [N· m/deg] | Orientation [deg] | Eccentricity [-] |
| --- | --- | --- | --- |
| Complete | 28.68 (4.74) | 59.98 (17.44) | 0.83 (0.10) |
| AG1 | 17.78 (5.39) | 65.81 (12.87) | 0.92 (0.05) |
| AG2 | 10.01 (10.73) | -3.16 (10.87) | 0.89 (0.41) |

A significant linear relationship between GAL and *H* (*R*^2^ = 0.54, *F*_1,12703891_ = 1.50 · 10^7^, *p <* 0.001) was established via linear regression (Fig. 7, top), while a non - significant association between GAL and the mean of the binned stiffness orientation was observed (*R*^2^ = 0.55, *F*_1,6_ = 6.12, *p* = 0.06) (Fig. 7, bottom). However, a significant correlation between GAL and the variability of the joint stiffness orientation was established, with a smaller standard deviation within bins measured at greater GAL (*R*^2^ = 0.86, *F*_1,6_ = 39.09, *p* = 0.002). Moreover, within the set of admissible wrist joint stiffness we found 802,356 unique values of joint stiffness orientation within a tolerance of 10^−4^ deg, spanning the entire range [- 90, + 90) deg with the largest distance measured between two consecutive values of orientation equal to 0.01 deg. As such, we considered that all possible values of joint stiffness orientation are achievable in the neutral position with an appropriate muscle activation vector **a**.

**Fig. 7.**
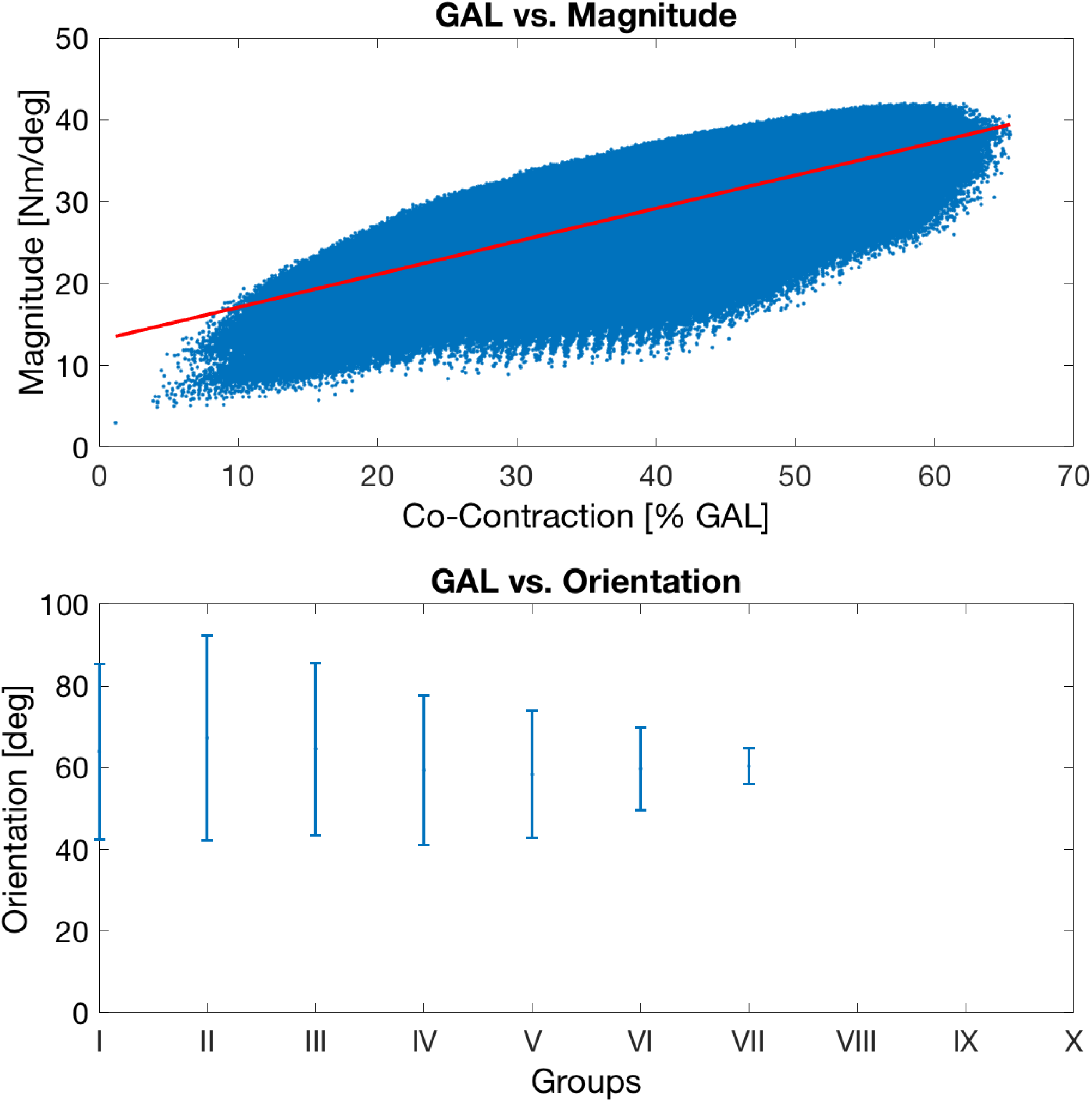
(Top) The plot represents the relationship between the magnitude of the achievable joint stiffness and the GAL; the red line is the fit obtained from linear regression. (Bottom) The plot represents the relationship between the orientation of the achievable joint stiffness and ten equally spaced group of GAL; the bars show the variability within each group.

### 3.3 Effect of different muscle groups

The systematic search for co - contraction of wrist and finger muscles returned 25,167 and 751,079 compatible activation vectors, respectively, which correspond to 4% and 13% of the tested activation vectors. Fig. 6 (center and right) shows the two sets of achievable stiffness color - coded by GAL. The distributions of the stiffness metrics for both virtual models are graphically represented in Fig. 8 with the mean and standard deviation reported in Tab. 3.

**Fig. 8.**
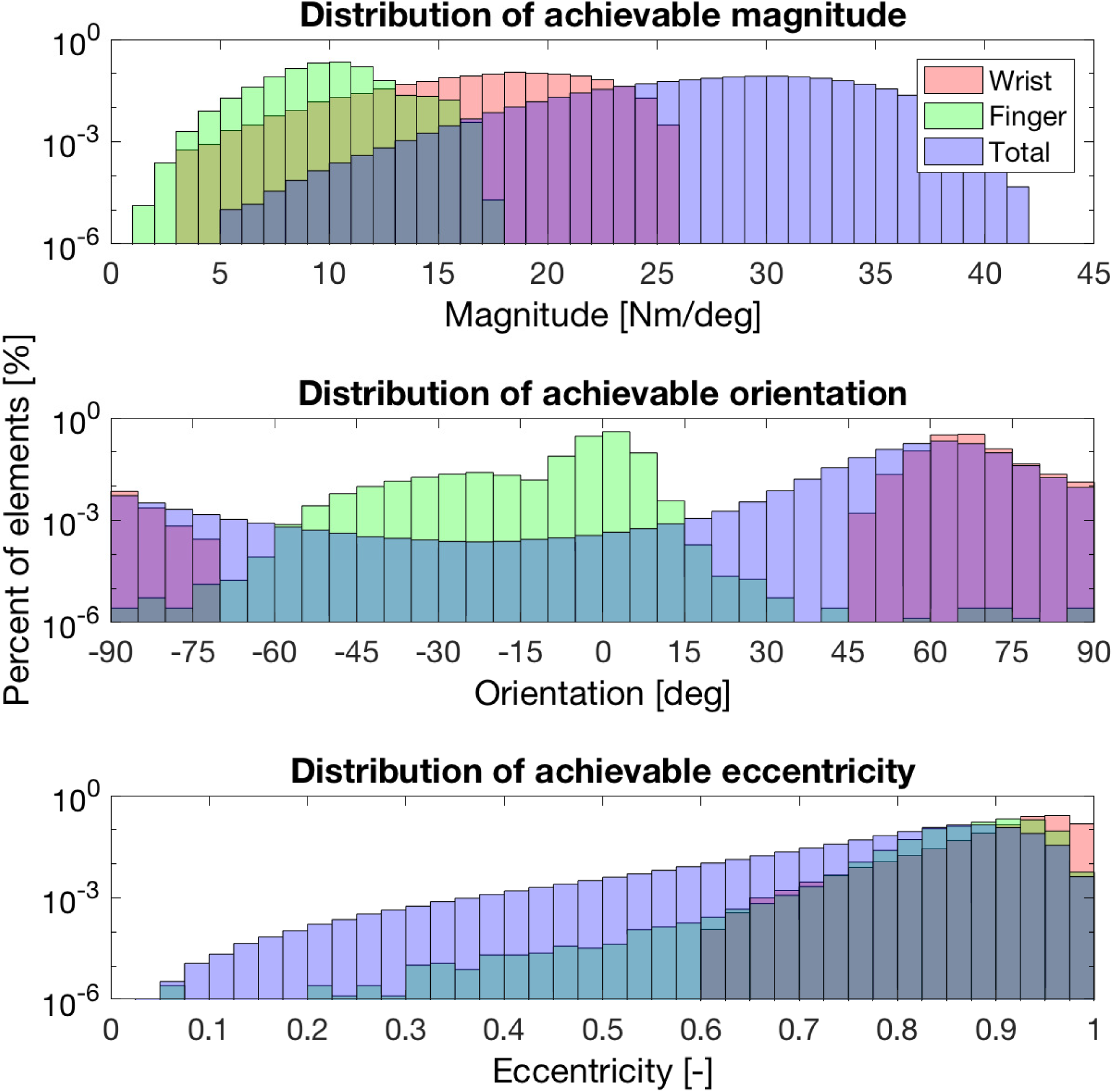
The histograms represent the distribution of the achievable magnitude (top) and orientation (center), and eccentricity (bottom), when different groups of muscles co - contract. Please note the logarithmic scale used for the vertical axis, which allows to appreciate the presence of a small number of feasible solutions for several values of magnitude, orientation, and eccentricity.

The test statistic rejected the null hypothesis that the mean of the distribution for AG1 and AG2 are equal for all the selected stiffness metrics — magnitude (*t*_13454969_ = 3.39 · 10^3^, *p <* 0.001), orientation (*t*_13454969_) = 2.59 · 10^3^, *p <* 0.001), and eccentricity (*t*_13454969_ = −534.59*, p <* 0.001).

## 4 Discussion

The purpose of this study was to use a realistic musculoskeletal model of the upper limb to study how different muscles and their activation contribute to modulating the active and passive component of the stiffness of the wrist joint. To this purpose, we developed a computational framework capable of estimating separately the two components of joint stiffness by integrating the parameters that describe the mechanical behavior of the musculotendon units in the musculoskeletal model with a short - range stiffness model. Our analysis based on the developed framework validated the passive wrist joint stiffness against previous experimental measurements, studied the effect of different muscle groups on the wrist joint passive stiffness in the neutral position, and studied the variability of the passive wrist joint stiffness in a 2D domain of wrist joint angles. Moreover, our analysis studied the effect of muscle co - contraction on the active wrist joint stiffness, establishing the set of stiffness values that are achievable via co - contraction of all relevant muscles, and studied the effect of different muscle groups on the wrist joint active stiffness. To our knowledge, no studies had been done to analyze the effect of muscle co - contraction in the wrist joint and only few [11, 12] studied the passive joint stiffness in the neutral posture. Our work extends the previous work to multiple wrist postures and for an exhaustive sets of muscle activation values. Moreover, our analysis based on MSMs created virtual models that allow to uniquely address the effect of different muscle groups on the passive and active stiffness of the wrist joint.

From the analysis of the passive stiffness, it is possible to observe that the MSM returns an estimate of the wrist joint stiffness that, in the neutral posture, is comparable with the physiological measurements in terms of orientation and eccentricity of the stiffness ellipse (the difference is less than 15 deg for the orientation and less than 25% difference for the eccentricity). The 60% difference in the stiffness magnitude can be justified by the fact that no ligaments, skin, nor cartilage structures are included in the model. Such structures, being in parallel with the muscles, have an effect on the overall joint stiffness as such, their absence leads to lower stiffness magnitudes. Furthermore, the MSM describes an individual with standard anatomy [36]; thus to compare model estimates with experimental values, a scaling of the model to a specific subject population would be necessary.

A second notable result observable from the passive stiffness analysis is the important role that finger muscles have in determining the wrist joint stiffness. As shown in Fig. 3, the wrist muscles alone are not capable of reproducing the physiological value of stiffness with errors of about 30 deg. However, the progressive inclusion of thumb (Fig. 3B, red line) and finger muscles (Fig. 3B, black line) produces a rotation of the ellipse and a decrease of the eccentricity bringing it to a condition comparable to physiological ones. The palm muscle is slack in the neutral posture, so its inclusion does not affect the passive joint stiffness.

Furthermore, we estimated for first time the passive joint stiffness throughout the wrist workspace and thus its variability in terms of magnitude, orientation and eccentricity. Our analysis shows a moderate variability for the magnitude (i.e. *c*_*v,H*_ = 0.39), a moderate - low variability for the orientation (*c*_*v*,Φ_ = 0.20), and a moderate - high variability for the eccentricity (*c*_*v,e*_ = 0.47). Moreover, it is interesting to observe that the orientation has a bi - modal distribution (Fig. 5) with the stiffness in the bottom right quadrant mostly characterized by an orientation centered about 90 deg and the remaining workspace characterized by an orientation centered about −60 deg.

Lastly, based on our framework, we investigated for the first time the effect of muscle co - contraction on the wrist joint stiffness in terms of orientation, eccentricity and magnitude of the stiffness ellipse. As shown in Fig. 6C and Fig. 7, muscle co - contraction produces two main effects on the joint stiffness: (*i*) it modulates the magnitude of the ellipses (Fig. 7A); (*ii*) it rotates the orientation of the principal semi - axis of the ellipse. Even though the fact that a greater muscle activation results in stronger co - contraction and therefore in increased joint stiffness has been already observed [16], the effect of co - contraction on the controllability of joint stiffness orientation was not fully understood. Our analysis shows that the variability of the orientation of the stiffness in the achievable set **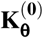** decreases with the increase in GAL. However, for low GAL and low stiffness magnitude, we measured feasible solutions almost for every value of stiffness orientation; as such, within this range of muscle co - contraction levels, there does not appear to be limitations on the set of achievable stiffness orientations. On the opposite, when the desired stiffness magnitude increases and thus the level of GAL, the variability of the orientation of the achievable set of stiffness values is lower, with admissible values clustered tightly around 60 deg.

By observing the distribution of the stiffness metrics, we finally investigated how co - contraction of different muscle groups contributes to modulating the wrist joint stiffness. Our analysis shows that the co - contraction of wrist and finger muscles produces achievable stiffness sets **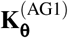** and **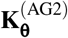** with different orientation, magnitude, and eccentricity. We relate the evidence that **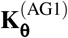** and **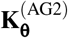** have statistically different orientation to the fact that each muscle may have a preferential direction in which it increases the stiffness when activated, a direction related to its moment arm. When only a group of muscles is allowed to co - contract the effect is that the stiffness increases mostly in the same direction, that may be different for different muscle groups. The difference in the stiffness magnitude mean for wrist and finger muscles is instead related to the fact that wrist muscles are weaker then wrist muscles, as such finger muscles have smaller influence on the magnitude of joint stiffness.

As an important methodological point, when studying the passive joint stiffness, is that there is no control over the applied joint torque, which is uniquely determined by the stretch of the passive fibers, directly resulting from the imposition of a wrist posture. As such, the presence of an equilibrium point, characterized by zero net torque, is not guaranteed. To calculate the wrist joint stiffness in each posture of the model, we assumed that the system would be in equilibrium with an external torque, equal and opposite to the one caused by the stretch of the passive fibers, applied in each posture of the workspace. As such, our analysis considered each point in the workspace as an equilibrium point, thereby allowing us to calculate the wrist joint stiffness for small deviations about that point. This procedure is the direct extension to a model - based analysis of the experimental techniques for measuring the stiffness of the wrist joint previously presented in [11–13].

Our study has two main limitations that results from our model - based approach. First, the model has unstable behavior in parts of the workspace for the passive condition; even though the unstable portion of the workspace decreases with the progressive inclusion of thumb and finger muscles, even when the entire set of muscles is considered, some areas of instability remain in wrist postures that are physiological. As such, since it is unclear whether the source of these instabilities is a reflection of real biomechanical properties or just a numerical problem [44], we reduced the workspace for stiffness calculation to a rectangular subset where the passive joint stiffness is stable. Moreover, the parameters of the SRS model have only been estimated for animal muscles [41]. Even though previous studies validated the use of this SRS model to describe experimental measurements of end - point stiffness measured during isometric contractions [32], accurate measurements of the γ parameter of the SRS model for human muscles are required to guarantee more reliable estimates of the joint stiffness.

## 5 Conclusion

In this study, we sought to gain a better understanding on the wrist joint stiffness and on its controllability by muscle co - contraction. We developed a novel computational framework based on a musculoskeletal model to estimate the active and passive components of joint stiffness.

Overall, the results of our study show that the thumb and finger muscles play a crucial role in determining the passive component of the wrist joint stiffness; in their absence, the model fails to reproduce the orientation of the wrist joint stiffness measured in prior experiments. Moreover, during active co - contraction, finger muscles produce a set of wrist joint stiffness that differs in magnitude and orientation with respect to the set obtained via co - contraction of the main wrist muscles. As such, when studying the wrist stiffness, it is of crucial importance to monitor the level of finger muscles activity in order to obtain reliable and repeatable measurements.

Furthermore, a second notable result is obtained when studying the controllability of the wrist joint stiffness with muscle co - contraction. Our analysis shows that co - contraction level positively correlates with the stiffness magnitude and negatively correlates with the variability in stiffness orientation. Our analysis showed that for low values of GAL, with appropriate selection of the activation vector, it is possible to modulate the orientation of the joint stiffness over 360 deg. This means that the range of possible orientations is potentially not bounded by biomechanical constraints; as such the limited capability in joint stiffness modulation observed in previous studies [26–28] may be caused primarily by neural constraints (e.g. muscle synergies).

## 6 Acknowledgments

We acknowledge support from the University of Delaware Research Foundation grant no. 16A01402, and from startup funds by the University of Delaware. The authors would like to thank Domenico Campolo for providing inspiring comments on preliminary versions of this work.

